# Detecting natural selection in trait-trait coevolution

**DOI:** 10.1101/2021.05.05.442737

**Authors:** Daohan Jiang, Jianzhi Zhang

**Affiliations:** Department of Ecology and Evolutionary Biology, University of Michigan, Ann Arbor, Michigan 48109, USA; Department of Quantitative and Computational Biology, University of Southern California, Los Angeles, California 90007, USA

**Keywords:** fly, morphology, mutation, modularity, pleiotropy, yeast

## Abstract

No phenotypic trait evolves independently of all other traits, but the cause of trait-trait coevolution is poorly understood. While the coevolution could arise simply from pleiotropic mutations that simultaneously affect the traits concerned, it could also result from multivariate natural selection favoring certain trait relationships. To gain a general mechanistic understanding of trait-trait coevolution, we examine the evolution of 220 cell morphology traits across 16 natural strains of the yeast *Saccharomyces cerevisiae* and the evolution of 24 wing morphology traits across 110 fly species of the family Drosophilidae, along with the variations of these traits among gene deletion or mutation accumulation lines (a.k.a. mutants). For numerous trait pairs, the phenotypic correlation among evolutionary lineages differs significantly from that among mutants. Specifically, we find hundreds of cases where the evolutionary correlation between traits is strengthened or reversed relative to the mutational correlation, which, according to our population genetic simulation, is likely caused by multivariate selection. Furthermore, we detect selection for enhanced modularity of the yeast traits analyzed. Together, these results demonstrate that trait-trait coevolution is shaped by natural selection and suggest that the pleiotropic structure of mutation is not optimal. Because the morphological traits analyzed here are chosen largely because of their measurability and thereby are not expected to be biased with regard to natural selection, our conclusion is likely general.

## BACKGROUND

Many phenotypic traits covary during evolution. For example, the logarithm of brain weight and that of body weight show a nearly perfect linear relationship across mammals [1, 2]. In theory, four processes may explain such trait-trait coevolution. First, it could arise simply from pleiotropic mutations that simultaneously influence these traits with a more or less constant ratio of effects [3-5], as has been previously shown empirically [6-10]. Second, trait covariation could arise from the linkage disequilibrium between genes controlling these traits [5, 11-13], but such trait covariation is expected to be restricted to closely related individuals due to the deterioration of linkage disequilibrium as a result of recombination. If the linkage disequilibrium is stably maintained due to, for example, chromosomal inversion, the involved linked genes can be regarded as a supergene with mutational pleiotropy [13]. For this reason, linkage disequilibrium is negligible except for trait covariation among closely related individuals. Third, shared ancestry can also create apparent trait correlations across lineages, which, however, can be explained away when the phylogenetic relationships are taken into account in correlation analysis [14]. Finally, trait covariation could be a result of natural selection for particular trait relationships that are advantageous, a phenomenon known as correlational selection or multivariate selection [2, 15-20].

Despite a long-standing interest in trait correlation in evolution [2, 13, 21], which is also referred to as phenotypic integration in the literature [22, 23], our understanding of the roles of mutation and selection in trait-trait coevolution remains limited. Most studies on the subject focused on a small number of traits that are physiologically or ecologically important [24], such as skull anatomy characters [25-30], behavioral syndrome (i.e., sets of correlated behavioral traits) [31, 32], and ecological or organismal traits correlated with the metabolic rate [33-37]; hence, they may not provide a general, unbiased picture of trait-trait coevolution. Additionally, it is the trait correlation resulting from standing genetic variation and its effect on adaptation that have received the most attention [38-44]. But, because standing genetic variation could have been influenced by selection [40], the resulting trait correlation may not inform the correlation produced by mutation. Not knowing the mutational correlation hinders a full understanding of the contribution of selection.

Related to trait-trait correlation is the concept of modularity. It has been hypothesized that it is beneficial for organisms to have a modular organization such that functionally related traits belonging to the same module covary and genotypes and/or phenotypes that lead to low fitness are less likely to occur [21, 25, 45-47]. Although modularity is a well-recognized feature of many trait correlation networks, the relative contribution of selection and mutational pleiotropy to modularity has not been assessed at the phenome scale [46-48].

To gain a general mechanistic understanding of trait-trait coevolution, we study the phenotypic correlations for a large number of trait pairs at the levels of mutation and long-term evolution; natural selection is inferred when the evolutionary correlation between traits cannot be fully explained by the mutational correlation. We also ask if the overall pattern of trait correlation (i.e., phenotypic integration) differ at the two levels. Our primary data include 220 cell morphology traits of the budding yeast *Saccharomyces cerevisiae* that have been measured in 4817 single-gene deletion lines [49], 89 mutation accumulation (MA) lines (for a subset of 187 traits) [50], and 16 natural strains with clear phylogenetic relationships [49, 51]. These traits were quantified from fluorescent microscopic images of triple-stained cells and were originally chosen for study because of their measurability regardless of potential roles in evolution and adaptation [49]. Subsequent studies found that these cell morphological traits are correlated with the yeast mitotic growth rate (i.e., a proxy for fitness) to varying degrees [7]. Hence, these traits may be considered representatives of phenotypic traits that have different contributions to fitness. Previous analyses of these traits among natural strains unveiled signals of positive selection on individual traits [52], but their potential coevolution has not been studied. While studying these trait pairs can offer a general picture of trait-trait coevolution, we recognize that the selective agent would be hard to identify should selection be detected, because the biological functions of these traits (other than correlations with the growth rate) are generally unknown [52]. To verify the generality of the findings from the yeast traits, we analyze another dataset that includes 12 landmark vein intersections on the fly wings that have been measured in 150 MA lines of *Drosophila melanogaster* [9] and 110 Drosophilid species [53]. At last, using computer simulations, we demonstrate how certain regimes of selection could explain the observed differences between mutational and evolutionary correlations.

## RESULTS

### Evolutionary correlations differ from mutational correlations for many trait pairs

To investigate if trait correlations in evolution can be fully accounted for by the correlations generated by mutation, we examined all pairs of the 220 yeast cell morphology traits previously measured. For each pair of traits, we computed the mutational correlation *COR*_M_, defined as Pearson’s correlation coefficient across 4,817 gene deletion lines (upper triangle in **Fig. 1A, Data S1**), and evolutionary correlation *COR*_E_, defined as Pearson’s correlation coefficient across 16 natural strains (lower triangle in **Fig. 1A, Data S1**) with their phylogenetic relationships (**Fig. S1**) taken into account (see Materials and Methods). Note that the original data contained 37 natural strains [51], of which 21 belong to the “mosaic” group [54, 55]—their phylogenetic relationships with other *S. cereviase* strains vary among genomic regions—so cannot be included in our analysis that requires considering phylogenetic relationships.

**Figure 1.**
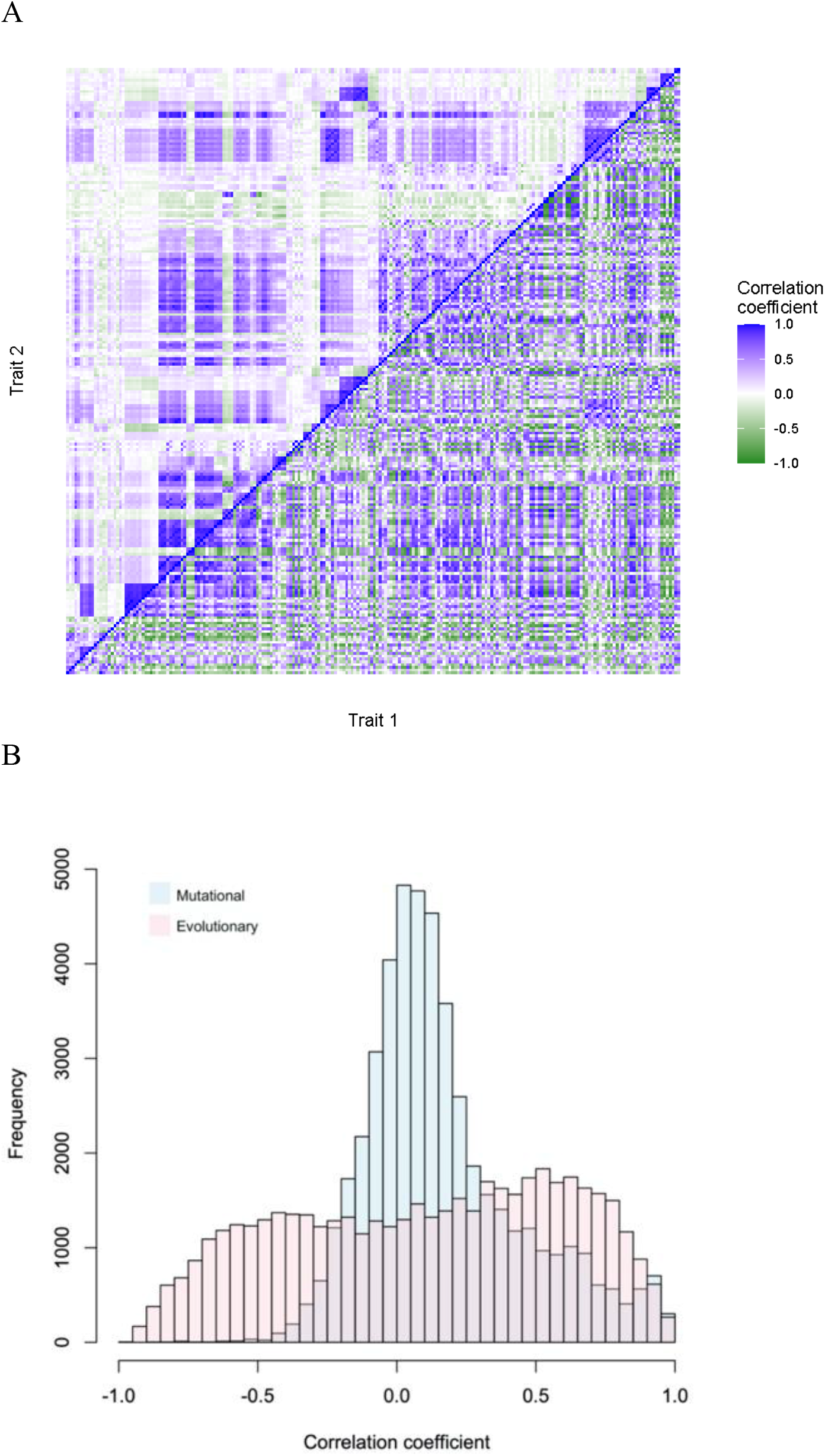
Mutational (*COR*_M_) and evolutionary (*COR*_E_) correlations for all pairs of the 220 yeast morphological traits. *COR*_M_ is based on yeast gene deletion lines. (A) *COR*_M_ (upper triangle) and *COR*_E_ (lower triangle) for all pairs of traits ordered according to their IDs. (B) Frequency distributions of *COR*_M_ and *COR*_E_ across all trait pairs. The two distributions are significantly different (*P* < 10^−10^, Kolmogorov–Smirnov test).

We found that the frequency distribution of *COR*_E_ across all trait pairs differs significantly from that of *COR*_M_ (**Fig. 1B**), suggesting the action of selection. For each pair of traits, we transformed the *COR*_M_ and *COR*_E_ to *Z*-scores using Fisher’s *r*-to-*Z* transformation and conducted *Z*-test to determine whether the two correlations are significantly different. Of the 24,090 trait pairs examined, 6743 pairs (or 28.0%) have a *COR*_E_ that deviates significantly from *COR*_M_ at the false discovery rate (FDR) of 5% (**Table 1, Data S1**), suggesting that natural selection has shaped the coevolution of many trait pairs. To investigate whether the above result is biased because of the use of each trait in many trait pairs, we randomly arranged the 220 traits into 110 non-overlapping pairs and counted the number of pairs with *COR*_E_ significantly different from *COR*_M_. This was repeated 1,000 times to yield 1,000 estimates of the proportion of trait pairs with significantly different *COR*_E_ and *COR*_M_. The middle 95% of these estimates ranged from 14.5% to 40.1%, with the median estimate being 28.2%, almost identical to the result (28.0%) from all pairwise comparisons. Hence, there is no indication that using overlapping trait pairs has biased the estimate of the fraction of trait pairs with significantly different *COR*_E_ and *COR*_M_.

**Table 1.** Numbers of trait pairs with significantly different *COR*_E_ and *COR*_M_ in the yeast and fly data.

|  | Yeast |  |  |  |  |  | Fly (276 trait pairs) |  |  |
| --- | --- | --- | --- | --- | --- | --- | --- | --- | --- |
| | $COR_M$ from gene deletion lines (24,090 trait pairs) | | | $COR_M$ from MA lines (17,391 trait pairs) | | | | | |
|  | Observed<sup>a</sup> | Expected<sup>b</sup> | Fraction<sup>c</sup> | Observed | Expected | Fraction | Observed | Expected | Fraction |
| Strengthened | 2727 | 145 | 3% | 2395 | 17 | 3.5% | 68 | 0 | 0% |
| Weakened | 1221 | 78 | 1.4% | 1348 | 6 | 0.1% | 48 | 0 | 0% |
| Reversed | 2795 | 51 | 0% | 1403 | 2 | 0% | 28 | 0 | 0% |
| Total | 6743 | 302.5 | 0.7% | 5146 | 31 | 0.7% | 144 | 0 | 0% |

<sup>a</sup>Number of trait pairs with significantly different $COR_M$ and $COR_E$ inferred by a Z-test.
<sup>b</sup>Number of trait pairs that show a significant difference between $COR_M$ and $COR_E$ derived from neutral Brownian motion simulations. The median from 1000 simulations is shown.
<sup>c</sup>Fraction of the 1000 Brownian motion simulations where the number of trait pairs with significantly different $COR_M$ and $COR_E$ exceeds the number under the “observed” column.

To further test selection, we simulated neutral evolution along the yeast tree 1000 times under a Brownian motion model with the observed mutational covariance matrix *M* used as the mutational input, generating 1,000 simulated datasets. Before the simulation, we confirmed that the sampling error of our estimated *M* is negligible, likely because of the large number of mutants used in *M* estimation (**Table S1**; see Materials and Methods). From each simulated dataset, we calculated the number of trait pairs with *COR*_E_ significantly different from *COR*_M_. Only in 0.7% of the simulated data did we find this number equal to or greater than that from the actual data (**Table 1**), indicating that the observed evolutionary correlations between traits cannot be explained by the neutral Brownian motion model. The distribution of mutational effects can be asymmetric and skewed [56] while it is assumed normal in the Brownian motion model. Nevertheless, simulations showed that mutational bias will not render *COR*_E_ deviate from *COR*_M_ in the absence of selection and will not enlarge the variance of *COR*_E_ (**Table S2**; see Materials and Methods).

We divided the 6743 cases of significantly different *COR*_E_ and *COR*_M_ into three categories. In the first category, the trait correlation generated by mutation is strengthened by natural selection during evolution. A total of 2,727 trait pairs are considered to belong to this “strengthened” category (**Table 1**) because they satisfy the following criteria: *COR*_E_ and *COR*_M_ have the same sign and |*COR*_*E*_| > |*COR*_*M*_|, or *COR*_E_ and *COR*_M_ have different signs but only *COR*_E_ is significantly different from 0 (at the nominal *P*-value of 0.05) (**Fig. 2A**). In the second category, the trait correlation generated by mutation is weakened by natural selection during evolution. A total of 1,221 trait pairs satisfying the following criteria are classified into this “weakened” category (**Table 1**): *COR*_E_ and *COR*_M_ have the same sign and |*COR*_*E*_| < |*COR*_*M*_|, or *COR*_E_ and *COR*_M_ have different signs but only *COR*_M_ is significantly different from 0 (**Fig. 2B**). In the last category, the trait correlation generated by mutation is reversed in sign by natural selection during evolution. A total of 2,795 trait pairs satisfying the following criteria are in this “reversed” category (**Table 1**): *COR*_E_ and *COR*_M_ have different signs and are both significantly different from 0 (**Fig. 2C**).

**Figure 2.**
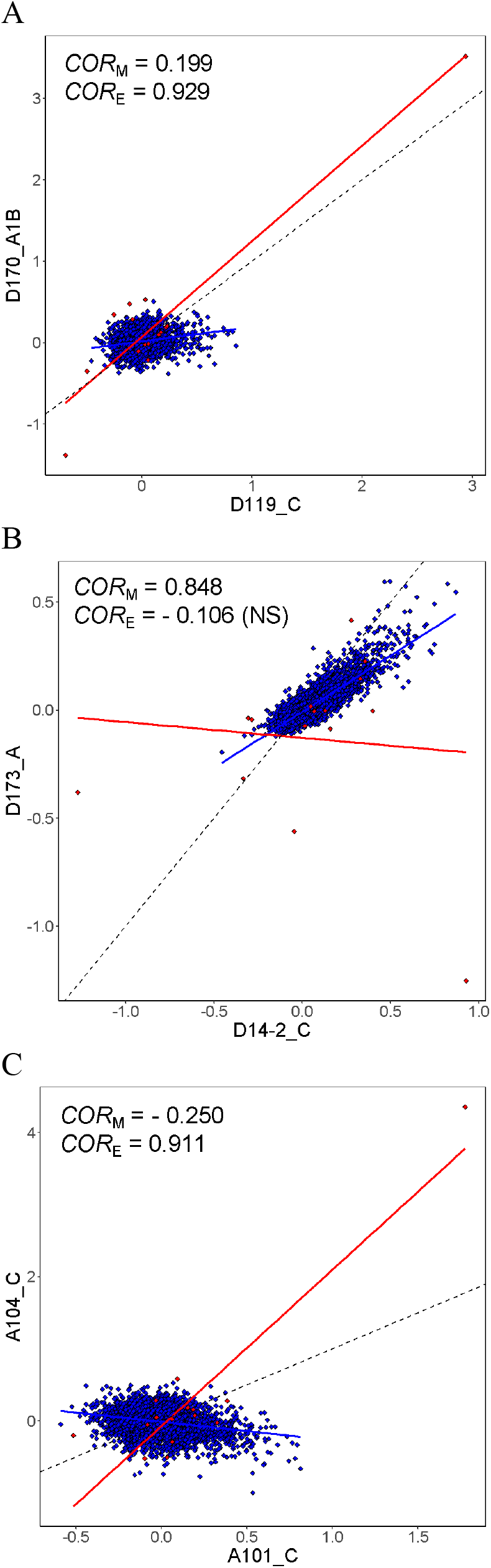
Examples of yeast trait pairs with *COR*_E_ significantly different from the corresponding *COR*_M_. (A) An example of evolutionarily strengthened correlation. (B) An example of evolutionarily weakened correlation. (C) An example of evolutionarily reversed correlation. Each blue dot represents a gene deletion line (a.k.a. mutant) while each red dot represents an independent contrast derived from natural strains. Blue and red lines are linear regressions between the standardized values of the two traits in mutants and independent contrasts, respectively, while the dotted blackline shows the diagonal (*y* = *x*). Trait IDs are shown along the axes. All *COR*_M_ and *COR*_E_ values shown are significantly different from 0 except when indicated by “NS” in the parentheses.

To assess the robustness of the selection signals detected, we repeated the above analysis using *COR*_M_ estimated from 89 mutation accumulation (MA) lines [43] (**Fig. S2A, Data S1**). Again, the overall frequency distribution across all trait pairs differs significantly between *COR*_*E*_ and *COR*_*M*_ (**Fig. S2B**). We found that 5,146 trait pairs exhibit a significantly different *COR*_E_ from the corresponding *COR*_*M*_ (**Table 1, Data S1**), supporting a role of selection in the coevolution of many trait pairs. When comparing the analysis using *COR*_M_ from gene deletion lines and that using *COR*_M_ from MA lines, we found 990 trait pairs to exhibit selection signals and fall into the same category in both analyses, including 275 pairs in the “strengthened” category, 223 pairs in the “weakened” category, and 574 pairs in the “reversed” category. All of these numbers substantially exceed the corresponding expected random overlaps (*P* < 0.001 based on 1,000 random draws in each case; the medians across the 1,000 draws are 271, 68 and 163, respectively), suggesting the reliability of both analyses. Although mutations in MA lines are more natural than those in gene deletion lines, the number of MA lines is much smaller than the number of gene deletion lines and only 187 of the original 220 traits were measured in the MA lines. For these reasons, we focused on the *COR*_M_ estimated from the gene deletion lines in subsequent analyses.

To examine the generality of the above yeast-based findings, we analyzed the 24 wing morphology traits of Drosophilid flies. The *COR*_M_ and *COR*_E_ have been previously estimated from 150 MA lines [9] and 110 Drosophilid species, respectively (**Fig. S3A, Data S1**). The overall frequency distribution across all trait pairs differs significantly between *COR*_*E*_ and *COR*_*M*_ (**Fig. S3B**). Of the 276 pairs of traits, 144 (52.2%) showed a significant difference between *COR*_E_ and *COR*_M_ (**Table 1, Data S1**), indicating widespread actions of selection in the coevolution of fly wing morphology traits.

Together, these results demonstrate that, for many trait pairs, mutational and evolutionary correlations between morphological traits are more different than expected under neutrality. This observation suggests an important role of selection in shaping the strength and/or direction of trait correlation in evolution.

### Effects of different selection regimes on trait-trait coevolution

The strengthened, weakened, and reversed trait correlations in evolution may have resulted from different selection regimes. Below we consider various selection regimes that could potentially explain these types of difference between *COR*_M_ and *COR*_E_ (**Fig. 3**). First, when a specific allometric relationship between two traits is selectively favored, the population mean trait values are expected to be concentrated near the fitness ridge or the optimal allometric line, resulting in a strong evolutionary correlation between the traits (i.e., a high |*COR*_*E*_|) (**Fig. 3A**). Unless *COR*_M_ is already similar to *COR*_E_, we expect to see strengthened or reversed *COR*_E_ depending on *COR*_M_. Second, if there is a single fitness peak for an optimal combination of trait values and if there is sufficiently strong stabilizing selection on the optimal phenotype, the population mean phenotype should be restricted within a small range of the optimal phenotype in all directions in the phenotypic space regardless of the mutational variance. Consequently, *COR*_E_ is expected to be close to 0, which could account for a weakened evolutionary correlation relative to the mutational correlation (**Fig. 3B**). Finally, if the fitness optimum varies across lineages in a random fashion, the steady-state *COR*_E_ will be close to zero, potentially leading to the weakening of the evolutionary correlation relative to the mutational correlation (**Fig. 3C**).

**Figure 3.**
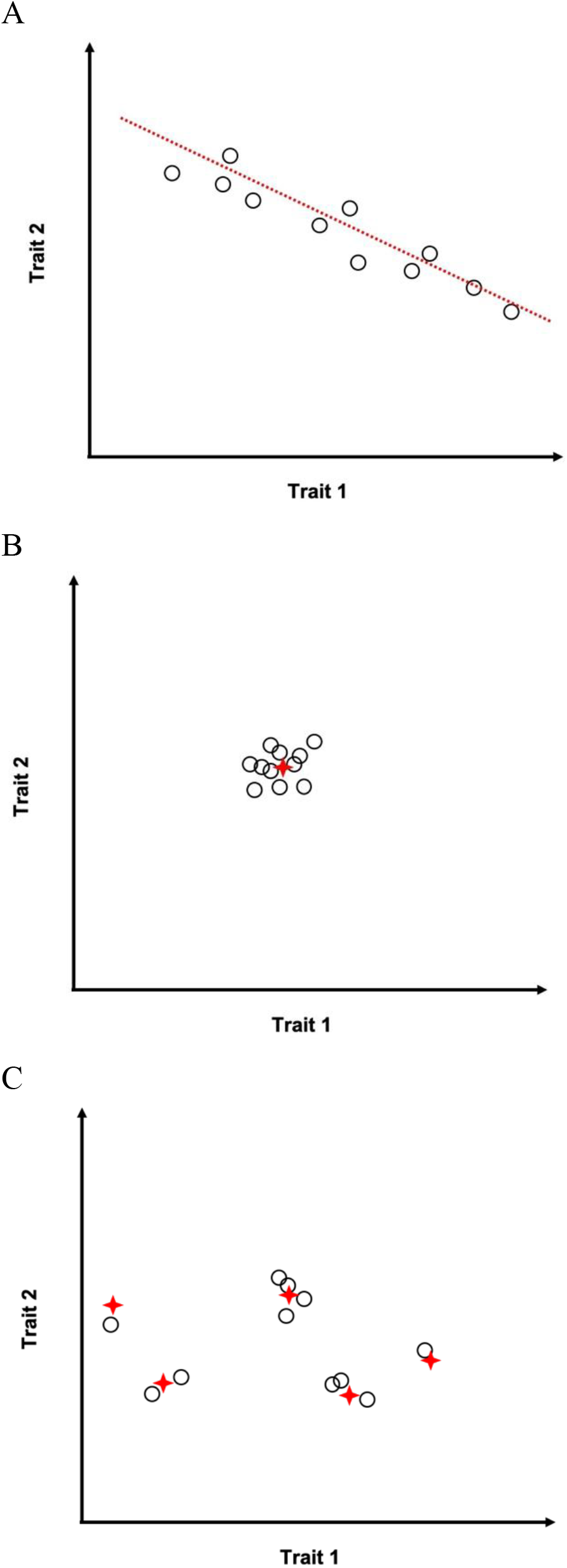
Schematic illustration of predictions made by models of trait-trait coevolution. Each circle represents the equilibrium mean phenotype of two hypothetical traits (trait 1 and trait 2) of a diverging lineage. (A) When a specific allometric relationship is selectively favored, the population mean phenotypes are distributed along the fitness ridge (i.e., the optimal allometric line shown in red), resulting in a strong trait correlation across lineages. (B) When a specific value is selectively favored for each trait, the population mean phenotypes are concentrated near the optimal phenotype (marked by the red cross) and the trait correlation across lineages is weak. (C) When different lineages have different optimal phenotypes (marked by red crosses) that are randomly distributed, the trait correlation across lineages is weak.

To verify these predictions, we simulated the evolution of two traits. Under each parameter set, we simulated 50 independent replicate lineages and computed the correlation coefficient, or *COR*_E_, between the traits across the replicate lineages at the end of the simulated evolution. This was repeated 200 times to obtain an empirical distribution of *COR*_E_. To evaluate the difference between *COR*_M_ and *COR*_E_, we examined the location of *COR*_M_ in the distribution of *COR*_E_; a significant (*P* < 0.05) difference is inferred if *COR*_M_ is in the left or right 2.5% tail of the *COR*_E_ distribution.

As expected, in the absence of selection, the distribution of *COR*_E_ is centered around *COR*_M_ (first block in **Table 2**). When a specific allometric relationship is selectively favored, a high |*COR*_E_| always emerges regardless of the *COR*_M_ used, resulting in either strengthened or reversed evolutionary correlations (*P* < 0.005 for all parameter sets examined; the second to fifth blocks in **Table 2**). By contrast, stabilizing selection of an optimal phenotype leads to weakened correlation across replicate lineages when |*COR*_*M*_| is not small (sixth block in **Table 2**). Finally, when different lineages have different phenotypic optima that are randomly picked from the standard bivariate normal distribution, weakened evolutionary correlations are generally observed except when *COR*_M_ is close to zero (bottom block in **Table 2**). These results suggest that the strengthened and reversed evolutionary correlations of yeast and fly morphological traits are likely caused by selections of allometric relationships, while the weakened correlations are likely caused by selections of individual traits either when there is a single optimal phenotype or when the optimal phenotype randomly varies among lineages.

**Table 2.** Parameters and results of simulations of trait-trait coevolution.

| Optimum | $COR_M$ | Median $COR_E$ at the end of simulation | Fraction of simulations with $COR_E > COR_M$ | $COR_E$ compared with $COR_M$ |
| --- | --- | --- | --- | --- |
| No optimum | 0.9 | 0.900 | 49.5% | No difference |
|  | 0.5 | 0.495 | 47.5% | No difference |
|  | 0.1 | 0.113 | 55.5% | No difference |
| $y = x^*$ | 0.9 | 1.000 | 100% | Strengthened |
|  | 0.5 | 1.000 | 100% | Strengthened |
|  | 0.1 | 1.000 | 100% | Strengthened |
| $y = 0.5x$ | 0.9 | 1.000 | 100% | Strengthened |
|  | 0.5 | 1.000 | 100% | Strengthened |
|  | 0.1 | 1.000 | 100% | Strengthened |
| $y = -0.5x$ | 0.9 | -0.995 | 0% | Reversed |
|  | 0.5 | -0.999 | 0% | Reversed |
|  | 0.1 | -1.000 | 0% | Reversed |
| $y = -x$ | 0.9 | -0.997 | 0% | Reversed |
|  | 0.5 | -0.999 | 0% | Reversed |
|  | 0.1 | -1.000 | 0% | Reversed |
| (0, 0) | 0.9 | 0.0213 | 0% | Weakened |
|  | 0.5 | 0.00142 | 0.5% | Weakened |
|  | 0.1 | -0.0109 | 24% | No difference |
| Drawn from $N(\mathbf{0}, \mathbf{1})$ | 0.9 | 0.0895 | 0% | Weakened |
|  | 0.5 | 0.0874 | 0% | Weakened |
|  | 0.1 | 0.0866 | 6% | No difference |

\*x and y respectively represent the values of the two traits considered.

### Selection for enhanced modularity of yeast morphological traits

While all of the above analyses focused on individual trait pairs, here we ask whether the overall trait correlation across divergent lineages is stronger or weaker than that created by mutation. As a measure of the overall level of trait correlation (i.e., overall integration), we calculated the variance of eigenvalues (*V*_eigen_) of the correlation matrix from divergent lineages and mutants, respectively. A greater *V*_eigen_ corresponds to a stronger overall correlation between traits because the eigenvalues become less evenly distributed as the absolute values of the correlation coefficients become larger [57]. However, the sample size (i.e., the number of strains) in the estimation of the correlation matrix also influences *V*_eigen_; a matrix estimated from a smaller sample naturally tends to have fewer positive eigenvalues and greater *V*_eigen_. To exclude the influence of this factor, we randomly sampled the mutant strains to obtain 5000 control datasets. Because the rank number of the evolutionary correlation matrix is 15 for the yeast data (i.e., 15 positive eigenvalues), each control dataset also consists of 15 randomly drawn strains such that the corresponding mutational correlation matrix also has 15 positive eigenvalues. We examined the location of the observed *V*_eigen_ in this distribution and computed a *P*-value based on this location (see Materials and Methods). For the yeast traits, *V*_eigen_ of the observed evolutionary correlation matrix exceeds that in 96% of control datasets (*P* = 0.08 in a two-tailed test; **Table 3**). Furthermore, only two of the 5000 control datasets have *V*_eigen_ significantly different from that of the observed evolutionary correlation matrix (Fligner-Kileen test). Hence, there is little evidence for a difference between the overall evolutionary correlation and the overall mutational correlation in yeast. For the fly data, the number of positive eigenvalues is unlimited by the sample size for both the evolutionary and mutational correlation matrices, hence we directly compared *V*_eigen_ between the two matrices, but found them to be similar (*P* = 0.459, Figner-Kileen test; **Table 3**). We also compared the overall integration between yeast and flies using *V*_eigen_/(*n*-1), where *n* is the number of traits examined. *V*_eigen_/(*n*-1) equals 0.157 and 0.268 for the yeast mutational and evolutionary matrices, respectively, whereas the corresponding values in flies are 0.153 and 0.190, respectively.

**Table 3.** Overall phenotypic integration (*V*_eigen_) and modularity (*CR*) at the levels of mutation and evolutionary divergence. Values at the level of mutation for yeast are medians of 1,000 control datasets. *P*-values for yeast are computed from locations of the observed values in the corresponding distributions of 5000 control datasets, while the *P*-value for fly is from a Fligner-Killeen test.

| Statistic | Taxon | Mutation | Divergence | $P$-value |
| --- | --- | --- | --- | --- |
| $V_{\text{eigen}}$ | Yeast | 34.414 | 58.656 | 0.0788 |
|  | Fly | 3.530 | 4.359 | 0.459 |
| $CR$ | Yeast | 0.649 | 0.997 | < 0.001 |

In addition to the overall level of trait correlation, we asked whether the correlational structure of traits exhibits different levels of modularity among divergent lineages when compared with that among mutants. To this end, we used a covariance ratio (*CR*) test [58] that compares covariance within and between pre-defined modules (see Materials and Methods). Specifically, we calculated *CR* for the evolutionary covariance matrix and compared it to the *CR* distribution based on 5000 mutational covariance matrices estimated from the randomly drawn subsets of mutants aforementioned. We treated the three non-overlapping categories of the yeast morphological traits—actin traits, nucleus traits, and cell wall traits [49]—as three modules (**Data S1**). We found that the *CR* of the evolutionary covariance matrix exceeded that of every control dataset (*P* < 0.001; **Table 3**), suggesting natural selection for increased modularity in evolution.

## DISCUSSION

By comparing the trait-trait correlation across mutants (*COR*_M_) with that across divergent lineages (*COR*_E_) for 24,090 pairs of yeast cell morphology traits and 276 pairs of fly wing morphology traits, we detected the action of natural selection in trait-trait coevolution. The fraction of trait pairs showing evidence for selection is substantially higher in the fly (52%) than yeast (28%) data (*P* < 10^−4^, chi-squared test). This is at least in part caused by a difference in statistical power, because the number of strains/species used for estimating *COR*_E_ is much greater for the fly (110) than yeast (16) data. It is likely that a higher fraction than 28% of the yeast trait pairs are subject to selection in their coevolution. Furthermore, our comparison between *COR*_E_ and *COR*_M_ intends to test selection on trait correlations common among the evolutionary lineages considered. If different evolutionary lineages have different trait correlations, the *COR*_E_ estimated from all lineages may not be significantly different from *COR*_M_ even when selection occurs in some or all of the lineages. In other words, our test is expected to underestimate the proportion of trait pairs subject to selection.

One potential biological explanation of the yeast-fly disparity in the prevalence of correlational selection is divergence time: the fly species represent a group that is tens of millions of years old while the yeast strains diverged from each other much more recently [53-55]. It is known that genetic correlations predict evolutionary correlations better over shorter timescales [38]. Similarly, selection might have had more time to decouple the pattern of evolutionary divergence from the mutational input in the flies but not yet in the yeast strains.

While we assumed that the mutants used carry all designed or natural mutations, extremely deleterious mutations such as lethal mutations are not represented. However, because such mutations are quickly selectively purged in natural populations, they should only be present transiently and are presumably unlikely to contribute to long-term evolution. Hence, their absence from our mutant data should not qualitatively alter our results.

We demonstrated by simulations that various selection regimes can explain differences between *COR*_M_ and *COR*_E_. In particular, strengthened or reversed *COR*_E_ relative to *COR*_M_ can occur when a specific allometric relationship is preferred, while weakened *COR*_E_ can occur under directional or stabilizing selection of individual traits. A notable difference between the simulation results and empirical observations is that the simulations tend to end up with extreme values of |*COR*_*E*_| (i.e., close to either 1 or 0) except in the case of neutrality, whereas the empirically observed |*COR*_*E*_| is usually less extreme even when *COR*_M_ and *COR*_E_ are significantly different. This is due to the fact that the simulation results usually represent steady-state correlations across lineages. That is, the mean phenotype of each lineage is at or near the corresponding optimum (if any); consequently, |*COR*_*E*_| is close to 1 when the optimum is a line and close to 0 when the optimum is a single combination of two trait values. However, the population mean phenotypes may not be close to their optima in some strains because of recent changes of the optima or the sparsity of mutations toward the optima, the latter of which is well known as a potential hindrance to adaptation [38, 42, 43, 59]. Another possibility is the existence of a wide range of preferred allometry such that there is no strong selection for extreme |*COR*_*E*_|. Finally, selection may not result in the preferred allometry between two traits because of the constraints from unconsidered traits [60].

It is worth noting that the yeast natural strains had been cultured in synthetic media before phenotyping [51] while the mutant strains were all grown in the rich medium YPD [49, 50]. Hence, it remains a possibility that the difference between *COR*_E_ and *COR*_M_ reported here contains a component caused by the environmental difference in phenotyping. Notwithstanding, our analysis suggests that this component is small (see Materials and Methods), which is expected because both media are meant to provide an ideal, stress-free environment for yeast growth. This said, future phenotyping in the same medium will be needed to validate our

While selection was detected for many trait pairs, a large fraction of trait pairs, especially in the yeast data, do not show a significant difference between *COR*_E_ and *COR*_M_. These trait pairs may be divided into two groups. In the first group, *COR*_E_ and *COR*_M_ are actually different, but the difference is not found significant due to the limited statistical power. As mentioned, we believe that a substantial fraction of yeast trait pairs belong to this category due to the relatively low statistical power in detecting the difference between *COR*_E_ and *COR*_M_ in the yeast data. In the second group, *COR*_E_ truly equals *COR*_M_, which could result from one of the following three scenarios. First, the specific trait-trait correlation does not impact fitness so evolves neutrally.

Second, the two traits have an intrinsic, immutable relationship (such as the hypothetical traits of body size and twice the body size), so will yield equal *COR*_E_ and *COR*_M_; this possibility can be tested by examining the correlation of the two traits across isogenic individuals that show non-heritable phenotypic variations [61]. The last and perhaps the most interesting scenario is that the trait-trait correlation impacts fitness and hence has driven the optimization of *COR*_M_ via a second-order selection [52, 59, 62, 63] such that the first-order selection of mutations that affect the two traits is no longer needed. However, the relative frequencies of these three scenarios are unknown.

In addition to pairwise trait correlations, we tested hypotheses regarding the evolution of overall phenotypic integration and modularity. In the yeast data, we observed a higher modularity across natural strains than across mutants but did not find evidence for a change of overall phenotypic integration in evolution. These results support the view of increasing modularity during evolution [21, 25, 45, 46, 64] but also suggest that modularity is enhanced by both strengthening trait-trait correlations within modules and weakening trait-trait correlations across modules. We found the overall integration lower for the fly than yeast traits, but whether this observation indicates a difference between different types of traits (i.e., cellular traits and multicellular organisms’ morphological traits) or between multicellular and unicellular organisms requires analyzing more species and traits.

Our analysis compared *COR*_M_ estimated from one yeast strain (BY) with *COR*_E_ estimated from 16 different strains, under the assumption of a constant *COR*_M_ across different strains. While it is a common practice to assume that the mutational architecture is more or less constant during evolution and to study phenotypic evolution by comparing mutational or genetic (co)variances in one species with those among different species [53, 65, 66], genetic variations affecting the genetic (co)variances of phenotypic traits have been reported [67-69]. As discussed earlier, such genetic variations may allow second-order selection of *COR*_M_. For instance, it has been hypothesized that the optimization of mutational (co)variances driven by selection for mutational robustness and/or adaptability can lead to modularity [21, 46]. It has indeed been found in the study of *Drosophila* gene expression traits that variational modules identified from mutants can be predicted to some extent by functional grouping of genes (i.e., Gene Ontology terms), although there is still much difference between functional modules and modules resulting from mutational pleiotropy, suggesting that optimization of the mutational architecture is far from complete even if it did take place [47]. Even without second-order selection, *COR*_M_ could still vary across strains because the pleiotropic effects of a mutation can vary by the environment and genetic background [19, 70, 71]. Regardless, in the future, it would be desirable to measure mutant phenotypes from multiple lineages to investigate whether *COR*_M_ evolves, how rapidly it evolves, and whether its evolution is largely neutral or adaptive.

Our analysis of the yeast dataset is subject to a major limitation resulting from the structure of the dataset. As many yeast strains are mosaic, only a small number of strains (16) were used in our study. Most of the remaining strains fall in one clade (**Fig. S1**), which is the Wine/European clade [54, 55]. That is, a substantial fraction of evolution along the yeast tree took place on internal branch(es), which would further reduce the effective sample size [72]. As a result, the *COR*_E_ estimate may not be very accurate, and the selection test suffers from low statistical power. It would be desirable if more non-mosaic strains from non-Wine/European clades are included. Another caveat regarding the calculation of *COR*_E_ is that correction methods like independent contrast do not always sufficiently account for the tree structure and can be susceptible to singular evolutionary events (e.g., shift of evolutionary rate in a clade) [73]; in our case, such a singular event could have taken place in the Wine/European clade after it had split from other yeast strains.

In summary, we detected the action of natural selection in shaping trait-trait coevolution. Because the traits analyzed here, especially the yeast traits, were chosen almost exclusively due to their measurability, our results likely reflect a general picture of trait-trait coevolution. Measuring these yeast traits in additional divergent natural strains with clear phylogenetic positions could improve the statistical power and clarify whether the fraction of trait pairs whose coevolution is shaped by selection is much greater than detected here. Finally, the detection of selection for enhanced modularity of the yeast traits analyzed supports the hypothesis that modularity is beneficial [21, 25]. The detection of selection in trait-trait coevolution and selection for enhanced modularity suggests that the current pleiotropic structure of mutation is not optimal. This nonoptimality could be due to the weakness of the second-order selection on mutational structure and/or a high dependence of the optimal mutational structure on the environment, which presumably changes frequently. Future studies on how the mutational structure evolves will likely further enlighten the mechanism of trait-trait coevolution.

## CONCLUSION

In this study, we analyzed morphological traits of yeast and flies and compared patterns of trait-trait correlation at the levels of mutation and long-term evolution. In both datasets, we discover that the evolutionary correlation differs significantly from the mutational correlation for numerous trait pairs, revealing a role of natural selection in trait-trait coevolution. We also provide evidence for selection for enhanced modularity of the yeast traits. Insights gained in this study can be summarized as follows:

1. Can trait-trait correlations in long-term evolution be explained by mutations? Our analyses showed that some correlations observed across divergent lineages differ significantly from correlations created by mutations. In addition, the pattern of phenotypic covariance among natural yeast strains has stronger modularity (i.e., stronger within-module correlations and/or weaker between-module correlations) than among mutants. These observations together indicate that selection likely played a role in shaping trait correlations in long-term evolution.
2. What evolutionary forces drive trait-trait correlation during evolution? Our simulations show how various selection regimes render the pattern of correlation during evolution different from that caused by mutation. Some types of differences, including strengthening and reversal of correlations, are explained by selection for an optimal allometric relationship, but not selection on individual traits.

## MATERIALS AND METHODS

### Phenotypic data

The *S. cerevisiae* cell morphology traits were previously measured by analyzing fluorescent microscopic images. Three phenotypic datasets were compiled and analyzed in this study, including (i) 220 traits measured in 4,718 gene deletion lines that each lack an nonessential gene [49], (ii) the same 220 traits measured in 37 natural strains [51], and (iii) 187 of the 220 traits measured in 89 mutation accumulation (MA) lines [50]. When comparing patterns of trait correlation between two datasets, we used traits available in both datasets. For each deletion strain, many cells (95 on average) were phenotyped, and the average trait value of all these cells were used to represent the strain in our analyses.

Three types of traits were measured in the deletion strains and the natural strains, including actin traits (i.e., measurements based on dyed actin cytoskeleton), cell wall traits (i.e., measurements based on dyed mannoprotein and cell wall markers), and nucleus traits (i.e., measurements based on dyed nuclear DNA) [49, 51]. These three categories were treated as three modules in our analysis of modularity. Only the cell membrane traits and nucleus traits were measured in the MA lines [50].

Before the analyses, we first standardized all trait values by converting each trait value to the natural log of the ratio of the original trait value to a reference such that the distributions become approximately normal and suitable for the *Z*-test. The standardized value of the *i*th trait in the *j*th strain is 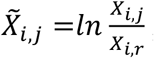, where *X*_*i,j*_ is the original trait value and *X*_*i,r*_ is the trait value of the reference. For the gene deletion lines, the reference is the wild-type BY strain. For the MA lines, the reference is the progenitor strain used in MA. For natural strains, the reference is the same as the reference of the mutant strains to be compared with (i.e., wild-type BY or progenitor of the MA lines).

The locations of 12 vein intersections on the fly wing were previously measured in 150 MA lines of *Drosophila melanogaster* and a mutational covariance matrix was estimated [9]. Because each intersection is described by two coordinates, which are counted as two traits, there are 24 traits in this dataset. These traits were also measured in 110 Drosophilid species and an evolutionary covariance matrix was estimated with species phylogeny taken into account [53]. Both matrices are based on log-scale trait values.

### Influence of the sampling error on the correlational structure

To evaluate the influence of sampling error on the estimated mutational covariance matrix (i.e., the *M* matrix) of yeast or fly, we took samples (vectors of phenotypes) from the multivariate distribution of *M* (4,817 samples for yeast gene deletion data and 150 samples for fly MA data), estimated a covariance matrix (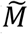) from these samples, and calculated Pearson’s correlation coefficient between the eigenvalues of *M* and 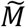.. For instance, for the yeast data, *M* and 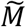 each has 220 eigenvalues, and we calculated the correlation between these two sets of eigenvalues as a measurement of similarity between *M* and 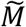.. This was repeated 1,000 times and the distribution of the correlation coefficient was used to evaluate the potential impact of sampling error on *M*.

### Impact of the environmental difference on the correlational structure of the yeast traits

Because the natural strains of yeast had been grown in synthetic media before phenotyping [51] while the mutant strains were all grown in the rich medium YPD [49, 50], we tested whether this environmental difference affected the correlational structure of the yeast morphological traits under consideration. Specifically, we examined whether the phenotype of the BY strain grown in synthetic media (referred to as “synthetic phenotype” for short) falls in the distribution of 123 biological replicates of BY grown in YPD (referred to as “YPD phenotypes” for short). The phenotypes were normalized in the way described earlier with the mean phenotype of the YPD replicates used as the reference. We decomposed YPD phenotypes into principal components (PCs) and focused on the first three PCs, which together explained 67.5% of the variance among the 123 YPD phenotypes. We then calculated the values of the three PC traits of the synthetic phenotype. The synthetic phenotype is in the central 95% of the distribution of the YPD phenotypes for each of the three PC traits, indicating a lack of major effect of the difference between synthetic and YPD media on the correlational structure of the yeast traits concerned.

### Comparison between mutational and evolutionary correlations

To take into account the phylogenetic relationships among yeast strains in estimating *COR*_E_, we utilized a distance-based tree previously inferred [55] (**Fig. S1**). Strains with mosaic origins inferred in the same study [55] were removed before analysis, resulting in 16 remaining natural strains. Because the BY strain was not included in the data file in that study [55], W303, a laboratory strain closely related to BY, was chosen to represent BY. We obtained the evolutionary covariance matrix using the *ratematrix* function from the R package *geiger* [74, 75], which calculates evolutionary covariances using the independent contrast method [14]. The evolutionary covariance matrix was then converted to the corresponding correlation matrix.

To test whether the observed pairwise trait correlation at the level of evolutionary divergence is significantly different from that expected by mutation alone for each pair of traits, we first converted both correlations to *Z*-scores by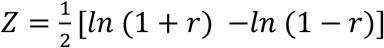, where *r* is the correlation coefficient. The testing statistic was then computed by 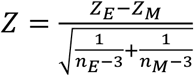, where *Z*_E_ and *Z*_M_ are *Z*-scores converted from *COR*_E_ and *COR*_M_, respectively, *n*_E_ is the number of independent contrasts, which equals the number of natural strains minus one, and *n*_M_ is the number of mutant strains. Two-sided *P*-value was calculated from each *Z* and converted to adjusted *P*-value following the Benjamini-Hochberg procedure [76]. An adjusted *P*-value below 0.05 indicates selection.

To see how many trait pairs would show a significant difference between *COR*_E_ and *COR*_M_ under neutrality, we simulated neutral evolution along the phylogenetic tree that had been used in estimating *COR*_E_. A Brownian motion model was used to simulate neutral phenotypic evolution such that the amount of evolution in branch *i* is *M*_*i*_*l*, where *M*_*i*_ is a vector sampled from the multivariate normal distribution of the mutational covariance matrix *M* and *l* is the branch length. Sampling was performed using the *rmvnorm* function in the R package *mvtnorm* [77]. The starting value of each trait is 0 in all simulations. The phenotypic value of each strain was obtained by adding up the amount of evolution on all branches ancestral to the strain. This was repeated 1,000 times to generate 1,000 datasets.

To account for the difference in *V*_eigen_ caused by different sample sizes in estimating the correlation matrices, we randomly sampled subsets of the gene deletion strains. Because the evolutionary correlation matrix has a rank number of 15 and has 15 positive eigenvalues, each subset consists of 15 strains randomly drawn from the 4718 gene deletion strains such that the mutational correlation matrix computed from each subset of mutants also has 15 positive eigenvalues. From each subset of strains, we computed *V*_eigen_, leading to a null distribution of *V*_eigen_. The observed *V*_eigen_ from the evolutionary correlation matrix is then compared with the null distribution; a significant difference is inferred if the observed value falls in either the left or right 2.5% tail.

To test whether there exists a significant modular structure among traits, we performed the covariance ratio (*CR*) test. For each pair of predefined modules, traits were first re-ordered such that traits belonging to each module were located in the upper-left and lower-right corners of the covariance matrix, respectively, and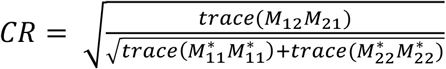, where *M*_12_ and *M*_21_ are the upper-right and lower-left sections of the original covariance matrix, respectively, containing all between-module covariances, 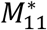 is the upper-left section with diagonal elements replaced by zeros, 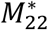 is the lower-right section with diagonal elements replaced by zeros, and *trace*(*M*) denotes the trace, or the sum of diagonal elements, of matrix *M* [58]. Because three modules were defined in the yeast data, the average of all pairwise *CR* values was used to represent the overall modularity. A test for selection on *CR* was performed following the test of selection on *V*_eigen_.

### Computer simulation of trait-trait coevolution under selection

In each simulation, we considered a pair of traits with equal amounts of mutational variance *V*_M_, which is set to be 0.01. The mutational covariance matrix is thus 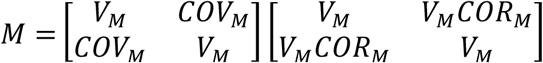, where *COV*_M_ is the mutational covariance. The number of mutations is a random Poisson variable with the mean equal to 1. The phenotypic effect of a mutation is drawn from the multivariate normal distribution of *M* using the *rmvnorm* function in the R package *mvtnorm* [77]. The starting phenotype is (0, 0) in all simulations.

We considered a Gaussian fitness function of 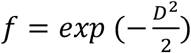 where *f* is the fitness and *D* is the distance between the current phenotype and the optimal phenotype. When there is a single fitness peak (i.e., the fitness optimum is a single point), *D* is the Euclidean distance defined by 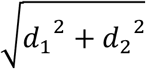, where *d*_1_ and *d*_2_ are the distances between the current phenotypic values of the two traits and their corresponding optima, respectively. When there is a fitness ridge (i.e., the fitness optimum is a line), *D* is the shortest distance from the current phenotype to the fitness ridge. The selection coefficient *s* equals 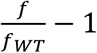, where *f* and *f*_WT_ are the fitness values of the mutant and wild type, respectively. The fixation probability of a newly arisen mutant is 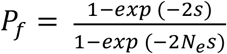 in a haploid population [78], where the effective population size *N*_e_ was set at 10^4^. After each unit time, the phenotypic effect of each mutation is added to the population mean with probability of *N*_*e*_*P*_*f*_; this probability is treated as 1 when *N*_*e*_*P*_*f*_ > 1 or when there is no selection as in the latter case 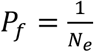. Combinations of parameters used in the simulations are listed in Table 2.

In simulations where different lineages were assigned different optima, each lineage’s optimum was obtained by independently drawing the optimal values of the two traits from the standard normal distribution. Before conducting simulations, we confirmed that the optima of the two traits are not correlated (correlation coefficient = 0.0882, *P* = 0.54, *t*-test).

### Computer simulation of trait-trait coevolution under mutational bias

To investigate the effect of mutational bias on trait correlation, we introduced the bias coefficient *B*. Each mutation, after being sampled from a multivariate normal distribution described above, was rescaled using *B*. Let the mutational effect be *m* = (*m*_1_, *m*_2_), where *m*_1_ and *m*_2_ are the effects on trait 1 and trait 2, respectively. The rescaled mutational effect, 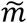, is obtained by

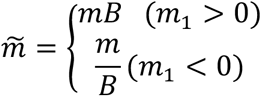

Because mutational effects are first drawn from a pre-set multivariate normal distribution and then rescaled, we examined if *COR*_M_ estimated from the rescaled effects (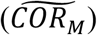) is different from the pre-set value of *COR*_M_. For each pre-set value of *COR*_M_, we obtained 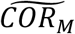 from 5,000 rescaled mutations. This was repeated 200 times with different random mutations, yielding 200 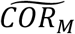 estimates. A series of different *B* values were used in the simulation (**Table S2**). For comparison, we also estimated *B* from yeast gene deletion lines and found the maximal *B* of any trait to be 1.503. To estimate *B* for a trait from the yeast gene deletion lines, we respectively calculated the mean trait value of all deletion lines with positive trait values and mean trait value of all deletion lines with negative values. We then computed the ratio of their absolute values with the greater absolute value used as the numerator. The square root of the ratio is *B*. We found that *COR*_M_ is always near the center of the distribution of these 200 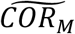 estimates (**Table S2**). Hence, mutational bias will not bias our test.

All analyses in this study were conducted in R [79].

## LIST OF ABBREVIATIONS

MA: mutation accumulation.
CR: covariance ratio.

## DECLARATION

### Ethics approval and consent to participate

Not applicable.

## Consent for publication

Not applicable.

## Availability of data and materials

The datasets generated and/or analysed during the current study are available at https://github.com/RexJiangEvoBio/Trait-Correlation.

## Authors’ contributions

D.J. and J.Z. designed the study. D.J. performed the analyses and prepared all figures. D.J. and J.Z. wrote the paper.

## Acknowledgements

We thank members of the Zhang lab and Dr. Matthew Pennell for valuable comments.

## REFERENCES

1. Gould SJ: Allometry and Size in Ontogeny and Phylogeny. Biol Rev 1966, 41(4):587–640.

2. Lande R: Quantitative Genetic Analysis of Multivariate Evolution, Applied to Brain - Body Size Allometry. Evolution 1979, 33(1):402–416.

3. Lande R: The Genetic Covariance between Characters Maintained by Pleiotropic Mutations. Genetics 1980, 94(1):203–215.

4. Wagner GP: Multivariate Mutation-Selection Balance with Constrained Pleiotropic Effects. Genetics 1989, 122(1):223–234.

5. Wagner GP, Zhang J: The pleiotropic structure of the genotype-phenotype map: the evolvability of complex organisms. Nature Reviews Genetics 2011, 12(3):204–213.

6. Dugand RJ, Aguirre JD, Hine E, Blows MW, McGuigan K: The contribution of mutation and selection to multivariate quantitative genetic variance in an outbred population of Drosophila serrata. Proc Natl Acad Sci U S A 2021, 118(31).

7. Ho WC, Zhang J: The genotype-phenotype map of yeast complex traits: basic parameters and the role of natural selection. Mol Biol Evol 2014, 31(6):1568–1580.

8. McGuigan K, Collet JM, McGraw EA, Ye YH, Allen SL, Chenoweth SF, Blows MW: The nature and extent of mutational pleiotropy in gene expression of male Drosophila serrata. Genetics 2014, 196(3):911–921.

9. Houle D, Fierst J: Properties of Spontaneous Mutational Variance and Covariance for Wing Size and Shape in Drosophila Melanogaster. Evolution 2013, 67(4):1116–1130.

10. McGuigan K, Collet JM, Allen SL, Chenoweth SF, Blows MW: Pleiotropic mutations are subject to strong stabilizing selection. Genetics 2014, 197(3):1051–1062.

11. Lande R: The maintenance of genetic variability by mutation in a polygenic character with linked loci (Reprinted). Genetics Research 2007, 89(5-6):373–387.

12. Gardner KM, Latta RG: Shared quantitative trait loci underlying the genetic correlation between continuous traits. Mol Ecol 2007, 16(20):4195–4209.

13. Saltz JB, Hessel FC, Kelly MW: Trait Correlations in the Genomics Era. Trends Ecol Evol 2017, 32(4):279–290.

14. Felsenstein J: Phylogenies and the Comparative Method. Am Nat 1985, 125(1):1–15.

15. Roff DA, Mostowy S, Fairbairn DJ: The evolution of trade-offs: Testing predictions on response to selection and environmental variation. Evolution 2002, 56(1):84–95.

16. Sinervo B, Svensson E: Correlational selection and the evolution of genomic architecture. Heredity (Edinb) 2002, 89(5):329–338.

17. Shoval O, Sheftel H, Shinar G, Hart Y, Ramote O, Mayo A, Dekel E, Kavanagh K, Alon U: Evolutionary trade-offs, Pareto optimality, and the geometry of phenotype space. Science 2012, 336(6085):1157–1160.

18. Bolstad GH, Cassara JA, Marquez E, Hansen TF, van der Linde K, Houle D, Pelabon C: Complex constraints on allometry revealed by artificial selection on the wing of Drosophila melanogaster. P Natl Acad Sci USA 2015, 112(43):13284–13289.

19. Svensson EI, Arnold SJ, Burger R, Csillery K, Draghi J, Henshaw JM, Jones AG, De Lisle S, Marques DA, McGuigan K et al: Correlational selection in the age of genomics. Nat Ecol Evol 2021.

20. Svensson EI: Multivariate selection and the making and breaking of mutational pleiotropy. Evol Ecol 2022, 36:807–828.

21. Wagner GP, Altenberg L: Perspective: Complex Adaptations and the Evolution of Evolvability. Evolution 1996, 50(3):967–976.

22. Olson EC, Miller RL: Morphological integration, Pbk. edn. Chicago: University of Chicago Press; 1999.

23. Pigliucci M: Phenotypic integration: studying the ecology and evolution of complex phenotypes. Ecol Lett 2003, 6(3):265–272.

24. Kingsolver JG, Hoekstra HE, Hoekstra JM, Berrigan D, Vignieri SN, Hill CE, Hoang A, Gibert P, Beerli P: The strength of phenotypic selection in natural populations. Am 653 Nat 2001, 157(3):245–261.

25. Goswami A, Smaers JB, Soligo C, Polly PD: The macroevolutionary consequences of phenotypic integration: from development to deep time. Philos Trans R Soc Lond B Biol Sci 2014, 369(1649):20130254.

26. Porto A, Sebastiao H, Pavan SE, VandeBerg JL, Marroig G, Cheverud JM: Rate of evolutionary change in cranial morphology of the marsupial genus Monodelphis is constrained by the availability of additive genetic variation. J Evol Biol 2015, 28(4):973–985.

27. Simon MN, Machado FA, Marroig G: High evolutionary constraints limited adaptive responses to past climate changes in toad skulls. Proc Biol Sci 2016, 283(1841).

28. Watanabe A, Fabre AC, Felice RN, Maisano JA, Muller J, Herrel A, Goswami A: Ecomorphological diversification in squamates from conserved pattern of cranial integration. Proc Natl Acad Sci U S A 2019, 116(29):14688–14697.

29. Fabre AC, Bardua C, Bon M, Clavel J, Felice RN, Streicher JW, Bonnel J, Stanley EL, Blackburn DC, Goswami A: Metamorphosis shapes cranial diversity and rate of evolution in salamanders. Nat Ecol Evol 2020, 4(8):1129–1140.

30. Navalon G, Marugan-Lobon J, Bright JA, Cooney CR, Rayfield EJ: The consequences of craniofacial integration for the adaptive radiations of Darwin’s finches and Hawaiian honeycreepers. Nature Ecology & Evolution 2020, 4(2):270−+.

31. Sih A, Bell A, Johnson JC: Behavioral syndromes: an ecological and evolutionary overview. Trends Ecol Evol 2004, 19(7):372–378.

32. Dochtermann NA, Dingemanse NJ: Behavioral syndromes as evolutionary constraints. Behav Ecol 2013, 24(4):806–811.

33. Martin RD: Relative brain size and basal metabolic rate in terrestrial vertebrates. Nature 1981, 293(5827):57–60.

34. Brown JH, Gillooly JF, Allen AP, Savage VM, West GB: Toward a metabolic theory of ecology. Ecology 2004, 85(7):1771–1789.

35. Glazier DS: A unifying explanation for diverse metabolic scaling in animals and plants. Biol Rev 2010, 85(1):111–138.

36. Pettersen AK, White CR, Marshall DJ: Metabolic rate covaries with fitness and the pace of the life history in the field. P Roy Soc B-Biol Sci 2016, 283(1831).

37. White CR, Marshall DJ, Alton LA, Arnold PA, Beaman JE, Bywater CL, Condon C, Crispin TS, Janetzki A, Pirtle E et al: The origin and maintenance of metabolic allometry in animals. Nat Ecol Evol 2019, 3(4):598–603.

38. Schluter D: Adaptive radiation along genetic lines of least resistance. Evolution 1996, 50(5):1766–1774.

39. Steppan SJ, Phillips PC, Houle D: Comparative quantitative genetics: evolution of the G matrix. Trends Ecol Evol 2002, 17(7):320–327.

40. Arnold SJ, Burger R, Hohenlohe PA, Ajie BC, Jones AG: Understanding the Evolution and Stability of the G-Matrix. Evolution 2008, 62(10):2451–2461.

41. Walsh B, Blows MW: Abundant Genetic Variation plus Strong Selection = Multivariate Genetic Constraints: A Geometric View of Adaptation. Annu Rev Ecol Evol S 2009, 40:41–59.

42. Agrawal AF, Stinchcombe JR: How much do genetic covariances alter the rate of adaptation? P Roy Soc B-Biol Sci 2009, 276(1659):1183–1191.

43. Blows MW, Mcguigan K: The distribution of genetic variance across phenotypic space and the response to selection. Mol Ecol 2015, 24(9):2056–2072.

44. Walter GM, Aguirre JD, Blows MW, Ortiz-Barrientos D: Evolution of Genetic Variance during Adaptive Radiation. Am Nat 2018, 191(4):E108–E128.

45. Wagner GP: A research programme for testing the biological homology concept. Novartis Found Symp 1999, 222:125–134; discussion 134-140.

46. Wagner GP, Pavlicev M, Cheverud JM: The road to modularity. Nature Reviews Genetics 2007, 8(12):921–931.

47. Collet JM, McGuigan K, Allen SL, Chenoweth SF, Blows MW: Mutational Pleiotropy and the Strength of Stabilizing Selection Within and Between Functional Modules of Gene Expression. Genetics 2018, 208(4):1601–1616.

48. Wang Z, Liao BY, Zhang JZ: Genomic patterns of pleiotropy and the evolution of complexity. P Natl Acad Sci USA 2010, 107(42):18034–18039.

49. Ohya Y, Sese J, Yukawa M, Sano F, Nakatani Y, Saito TL, Saka A, Fukuda T, Ishihara S, Oka S et al: High-dimensional and large-scale phenotyping of yeast mutants. P Natl Acad Sci USA 2005, 102(52):19015–19020.

50. Geiler-Samerotte KA, Zhu YO, Goulet BE, Hall DW, Siegal ML: Selection Transforms the Landscape of Genetic Variation Interacting with Hsp90. Plos Biol 2016, 14(10).

51. Yvert G, Ohnuki S, Nogami S, Imanaga Y, Fehrmann S, Schacherer J, Ohya Y: Single-cell phenomics reveals intra-species variation of phenotypic noise in yeast. BMC Syst Biol 2013, 7:54.

52. Ho WC, Ohya Y, Zhang JZ: Testing the neutral hypothesis of phenotypic evolution. P Natl Acad Sci USA 2017, 114(46):12219–12224.

53. Houle D, Bolstad GH, van der Linde K, Hansen TF: Mutation predicts 40 million years of fly wing evolution. Nature 2017, 548(7668):447−+.

54. Liti G, Carter DM, Moses AM, Warringer J, Parts L, James SA, Davey RP, Roberts IN, Burt A, Koufopanou V et al: Population genomics of domestic and wild yeasts. Nature 2009, 458(7236):337–341.

55. Peter J, De Chiara M, Friedrich A, Yue JX, Pflieger D, Bergstrom A, Sigwalt A, Barre B, Freel K, Llored A et al: Genome evolution across 1,011 Saccharomyces cerevisiae isolates. Nature 2018, 556(7701):339–344.

56. Hodgins-Davis A, Duveau F, Walker EA, Wittkopp PJ: Empirical measures of mutational effects define neutral models of regulatory evolution in Saccharomyces cerevisiae. Proc Natl Acad Sci U S A 2019, 116(42):21085–21093.

57. Pavlicev M, Cheverud JM, Wagner GP: Measuring Morphological Integration Using Eigenvalue Variance. Evol Biol 2009, 36(1):157–170.

58. Adams DC: Evaluating modularity in morphometric data: challenges with the RV coefficient and a new test measure. Methods Ecol Evol 2016, 7(5):565–572.

59. Hansen TF, Houle D: Measuring and comparing evolvability and constraint in multivariate characters. J Evolution Biol 2008, 21(5):1201–1219.

60. Houle D, Jones LT, Fortune R, Sztepanacz JL: Why does allometry evolve so slowly? Integr Comp Biol 2019, 59(5):1429–1440.

61. Geiler-Samerotte KA, Li S, Lazaris C, Taylor A, Ziv N, Ramjeawan C, Paaby AB, Siegal ML: Extent and context dependence of pleiotropy revealed by high-throughput single-cell phenotyping. Plos Biol 2020, 18(8).

62. Wagner A: Robustness and evolvability in living systems. Princeton, NJ: Princeton University Press; 2005.

63. Jones AG, Arnold SJ, Burger R: Evolution and stability of the G-matrix on a landscape with a moving optimum. Evolution 2004, 58(8):1639–1654.

64. Clune J, Mouret JB, Lipson H: The evolutionary origins of modularity. Proc Biol Sci 2013, 280(1755):20122863.

65. Ackermann RR, Cheverud JM: Detecting genetic drift versus selection in human evolution. Proc Natl Acad Sci U S A 2004, 101(52):17946–17951.

66. Lynch M: The Rate of Morphological Evolution in Mammals from the Standpoint of the Neutral Expectation. Am Nat 1990, 136(6):727–741.

67. Jerison ER, Kryazhimskiy S, Mitchell JK, Bloom JS, Kruglyak L, Desai MM: Genetic variation in adaptability and pleiotropy in budding yeast. Elife 2017, 6.

68. Jones AG, Burger R, Arnold SJ: Epistasis and natural selection shape the mutational architecture of complex traits. Nat Commun 2014, 5:3709.

69. Pavlicev M, Kenney-Hunt JP, Norgard EA, Roseman CC, Wolf JB, Cheverud JM: Genetic variation in pleiotropy: Differential epistasis as a source of variation in the allometric relationship between long bone lengths and body weight. Evolution 2008, 62(1):199–213.

70. Pavlicev M, Cheverud JM: Constraints Evolve: Context Dependency of Gene Effects Allows Evolution of Pleiotropy. Annual Review of Ecology, Evolution, and Systematics, Vol 46 2015, 46:413–434.

71. Wei X, Zhang J: Environment-dependent pleiotropic effects of mutations on the maximum growth rate r and carrying capacity K of population growth. Plos Biol 2019, 17(1):e3000121.

72. Ané C: Analysis of comparative data with hierarchical autocorrelation. Annals of Applied Statistics 2008, 2(3):1078–1102.

73. Uyeda JC, Zenil-Ferguson R, Pennell MW: Rethinking phylogenetic comparative methods. Syst Biol 2018, 67(6):1091–1109.

74. Pennell MW, Eastman JM, Slater GJ, Brown JW, Uyeda JC, FitzJohn RG, Alfaro ME, Harmon LJ: geiger v2.0: an expanded suite of methods for fitting macroevolutionary models to phylogenetic trees. Bioinformatics 2014, 30(15):2216–2218.

75. Revell LJ, Harmon LJ, Langerhans RB, Kolbe JJ: A phylogenetic approach to determining the importance of constraint on phenotypic evolution in the neotropical lizard Anolis cristatellus. Evol Ecol Res 2007, 9(2):261–282.

76. Benjamini Y, Hochberg Y: Controlling the False Discovery Rate - a Practical and Powerful Approach to Multiple Testing. J R Stat Soc B 1995, 57(1):289–300.

77. Genz AB, F.; Miwa, T.; Mi, X.; Leisch, F.; Scheipl, F.; Hothorn, T.: mvtnorm: Multivariate Normal and t Distributions. R package version 1.1-0, https://CRAN.R-project.org/package=mvtnorm. 2020.

78. Kimura M: On the probability of fixation of mutant genes in a population. Genetics 1962, 47:713–719.

79. R Core Development Team: R: A language and environment for statistical computing. 2010.

