## supplementary materials for "Detecting natural selection in trait-trait coevolution"

Supplementary materials for “Detecting natural selection in trait-trait coevolution” by D. Jiang & J. Zhang

The supplementary materials include:

Tables S1-S2

Legends of supplementary figures

Figures S1-S3

Data S1 (in a separate Excel file)

Table S1. Pearson’s correlation in eigenvalues between *M* and the covariance matrix estimated from a sample from *M* ($\tilde{M}$). Results from 1000 replications are shown.

| Taxon | Sample size | Minimum correlation coefficient | Median correlation coefficient |
| --- | --- | --- | --- |
| Yeast | 4,817 | 0.9993745 | 0.9999503 |
| Fly | 150 | 0.9698937 | 0.9967839 |

Table S2. Parameters and results of simulations of trait-trait coevolution in the presence of mutational bias.

| Mutational bias *B* | *COR*_M_ | Median $\tilde{{COR}_{M}}$ | Fraction of $\tilde{{COR}_{M}}>{COR}_{M}$ | Median *COR*_E_ at the end of simulation | Fraction of simulations with *COR*_E_ > *COR*_M_ | *COR*_E_ variance |
| --- | --- | --- | --- | --- | --- | --- |
| 2 | 0.9 | 0.900 | 45.5% | 0.885 | 31.5% | 0.0185 |
|  | 0.5 | 0.500 | 49.5% | 0.468 | 39.5% | 0.0175 |
|  | 0.1 | 0.102 | 54.5% | 0.0999 | 50% | 0.0210 |
| 10 | 0.9 | 0.900 | 50.5% | 0.863 | 13% | 0.0198 |
|  | 0.5 | 0.499 | 46.5% | 0.426 | 24.5% | 0.0261 |
|  | 0.1 | 0.101 | 54.5% | 0.0886 | 46.5% | 0.0168 |
| 100 | 0.9 | 0.900 | 52% | 0.861 | 12% | 0.0194 |
|  | 0.5 | 0.500 | 49% | 0.420 | 27% | 0.0261 |
|  | 0.1 | 0.101 | 52% | 0.0856 | 46.5% | 0.0211 |
| 1000 | 0.9 | 0.900 | 49.5% | 0.863 | 14% | 0.0218 |
|  | 0.5 | 0.500 | 49% | 0.443 | 30.5% | 0.0228 |
|  | 0.1 | 0.101 | 52.5% | 0.0849 | 44.5% | 0.0237 |

**Legends of supplementary figures**

**Fig. S1.** Neighbor-joining tree of the 16 natural yeast strains used in this study, based on 1,544,489 biallelic single nucleotide polymorphism (SNP) sites. Scale bar indicates genomic divergence level. The tree was based on the distance matrix downloaded from <http://1002genomes.u-strasbg.fr/files/1011DistanceMatrixBasedOnSNPs.tab.gz>. The inset at the top left coner shows the tree topology but the branch lengths are not drawn to scale.

**Fig. S2.** Mutational (*COR*_M_) and evolutionary (*COR*_E_) correlations for all pairs of the 187 yeast morphological traits. *COR*_M_ is based on yeast mutation accumulation lines. (A) *COR*_M_ (upper triangle) and *COR*_E_ (lower triangle) for all pairs of traits ordered according to their IDs. (B) Frequency distributions of *COR*_M_ (blue) and *COR*_E_ (red) across all trait pairs. The two distributions are significantly different (*P* < 10^-10^, Kolmogorov–Smirnov test).

**Fig. S3.** Mutational (*COR*_M_) and evolutionary (*COR*_E_) correlations for all pairs of the 24 fly wing morphological traits. (A) *COR*_M_ (upper triangle) and *COR*_E_ (lower triangle) for all pairs of traits ordered in the same way as in the original dataset. (B) Frequency distributions of *COR*_M_ (blue) and *COR*_E_ (red) across all trait pairs. The two distributions are significantly different (*P* = 0.0015, Kolmogorov–Smirnov test).

**Figure S1**

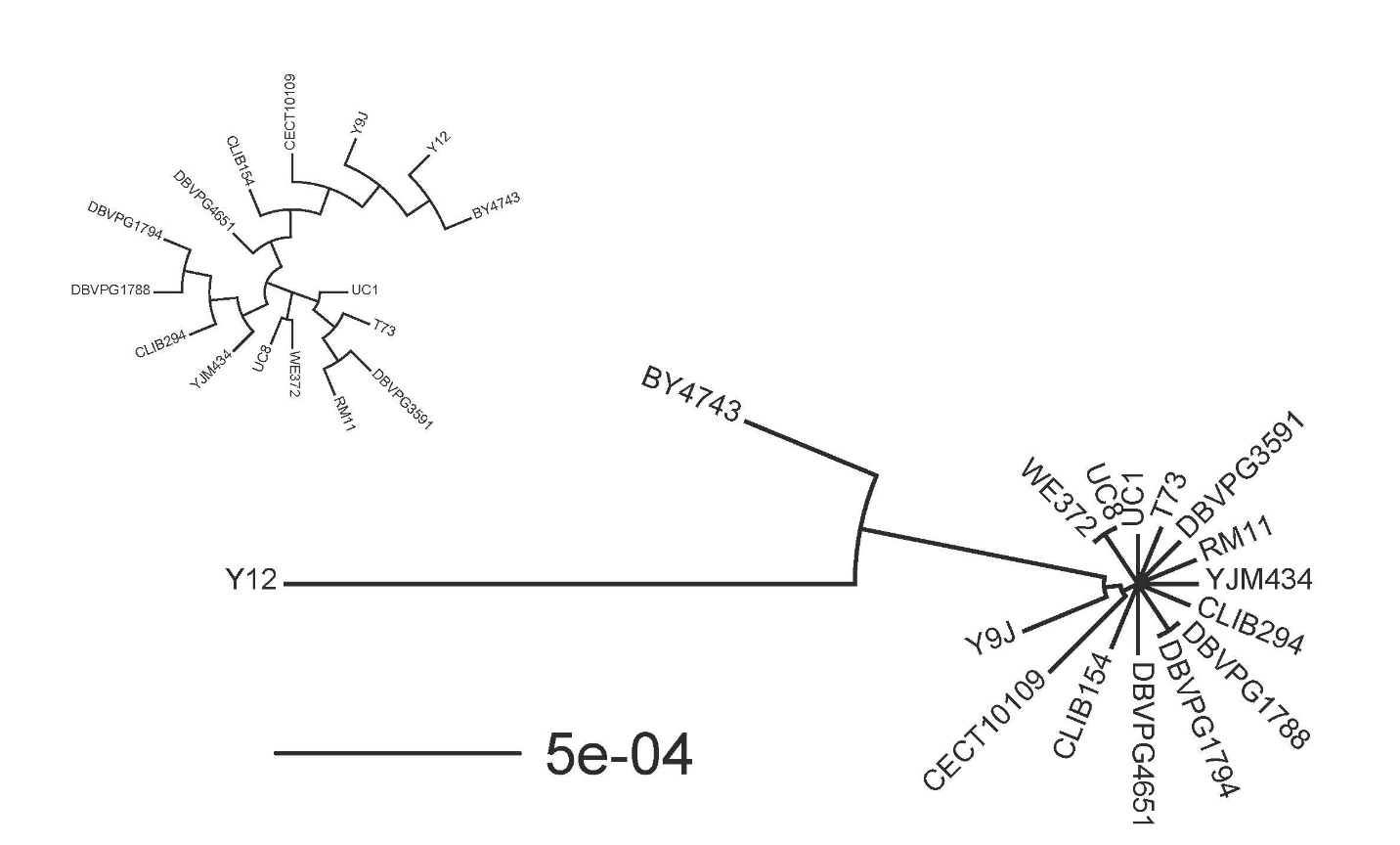

**Figure S2**

A

B
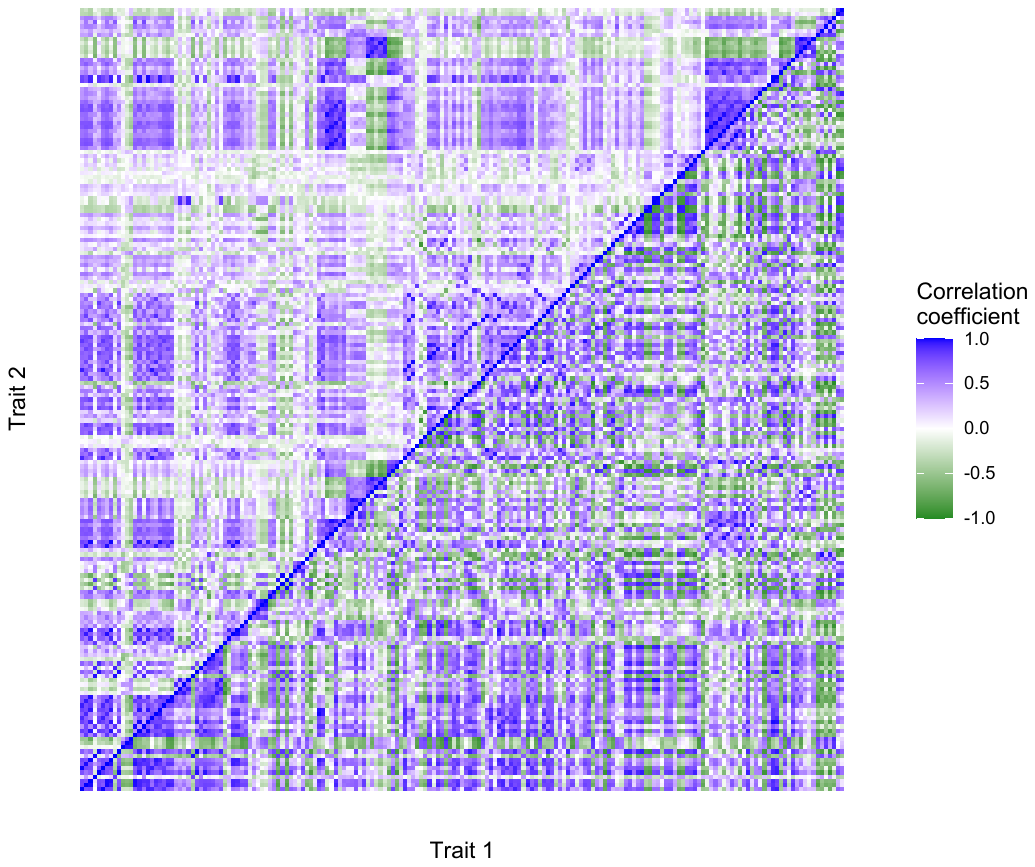

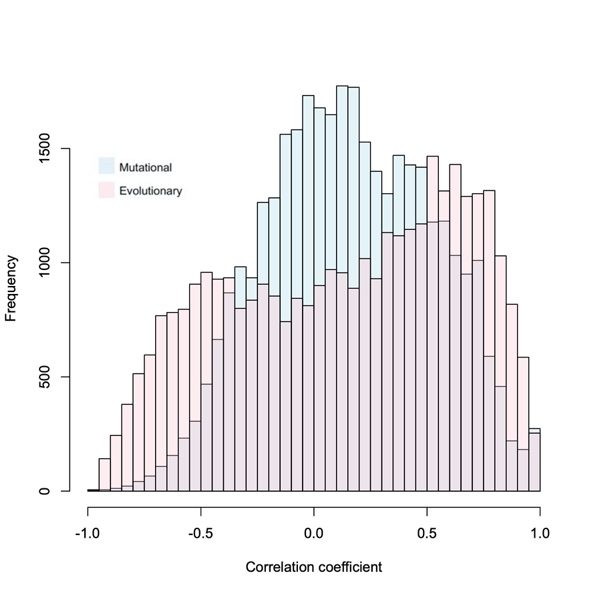

**Figure S3**

A

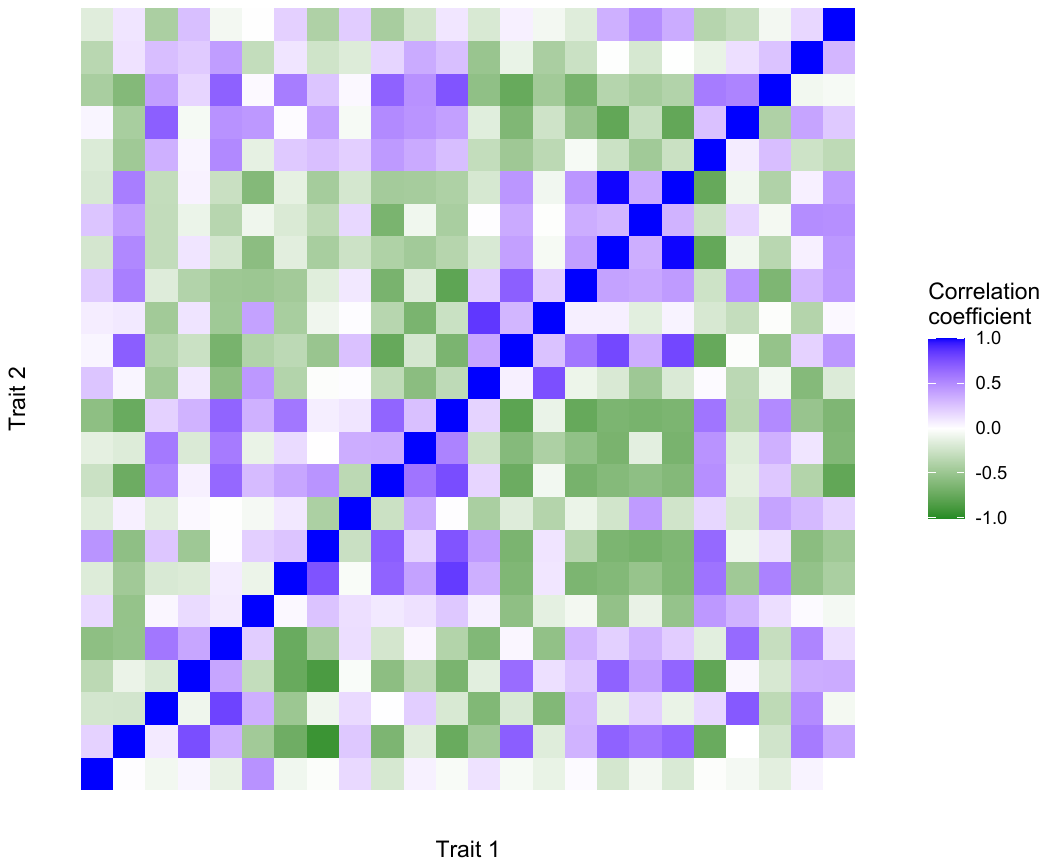

B

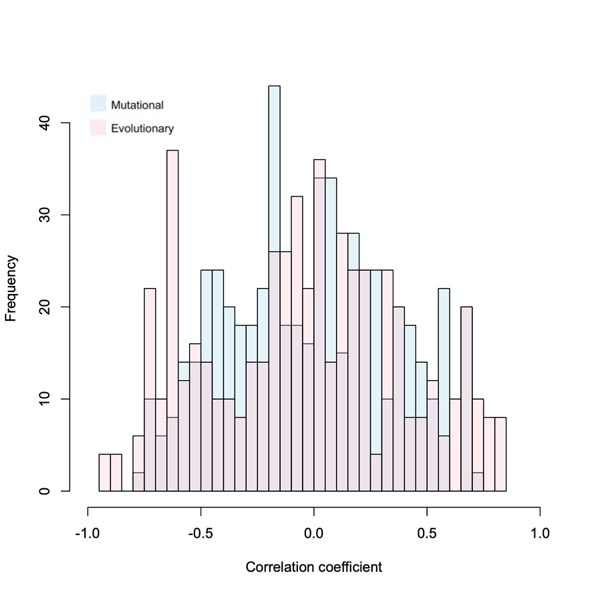
